# Transformation from independent to integrative coding of multi-object arrangements in human visual cortex

**DOI:** 10.1101/117432

**Authors:** Daniel Kaiser, Marius V. Peelen

**Affiliations:** Center for Mind/Brain Sciences, University of Trento, 38068 Rovereto (TN), Italy; Department of Education and Psychology, Freie Universität Berlin, 14195 Berlin-Dahlem, Germany; Donders Institute for Brain, Cognition and Behaviour, Radboud University, Nijmegen, The Netherlands

## Abstract

To optimize processing, the human visual system utilizes regularities present in naturalistic visual input. One of these regularities is the relative position of objects in a scene (e.g., a sofa in front of a television), with behavioral research showing that regularly positioned objects are easier to perceive and to remember. Here we use fMRI to test how positional regularities are encoded in the visual system. Participants viewed pairs of objects that formed minimalistic two-object scenes (e.g., a “living room” consisting of a sofa and television) presented in their regularly experienced spatial arrangement or in an irregular arrangement (with interchanged positions). Additionally, single objects were presented centrally and in isolation. Multi-voxel activity patterns evoked by the object pairs were modeled as the average of the response patterns evoked by the two single objects forming the pair. In two experiments, this approximation in object-selective cortex was significantly less accurate for the regularly than the irregularly positioned pairs, indicating integration of individual object representations. More detailed analysis revealed a transition from independent to integrative coding along the posterior-anterior axis of the visual cortex, with the independent component (but not the integrative component) being almost perfectly predicted by object selectivity across the visual hierarchy. These results reveal a transitional stage between individual object and multi-object coding in visual cortex, providing a possible neural correlate of efficient processing of regularly positioned objects in natural scenes.

## 1 Introduction

Our everyday environments are cluttered, consisting of a large number of separable objects. How does the visual system efficiently process such complex input?

Much of our knowledge about the neural mechanisms of object perception comes from neuroimaging studies using single objects as stimuli. These studies have linked object perception to activity in object-selective cortex (OSC), a region which preferentially responds to objects compared to scrambled objects (Grill-Spector, 2003; Malach et al., 1995) and exhibits reliably discriminable response patterns for different types of objects (Eger et al., 2008; Haxby et al., 2001; Kriegeskorte et al., 2008). Further neuroimaging work has demonstrated that OSC responses show some degree of size-and location-invariance (Cichy et al., 2011; Vuilleumier et al., 2002), partly reflect perceived rather than physical object properties (Haushofer et al., 2008; Kourtzi and Kanwisher, 2001), and are linked to behavioral performance in object recognition (Grill-Spector et al., 2000; Williams et al., 2007).

Recently, studies have moved towards more naturalistic conditions, measuring OSC responses when multiple objects are presented simultaneously. These studies have demonstrated that OSC codes multiple objects in an independent, linearly additive way: Responses to pairs of objects could be modeled as a linear combination (e.g., the average) of the responses to their constituent single objects (Agam et al., 2010; Baeck et al., 2013; Kaiser et al., 2014b; Kubilius et al., 2015; MacEvoy and Epstein, 2009; Reddy et al., 2009; Zoccolan et al., 2005), even when the objects were embedded in complex natural scenes (MacEvoy and Epstein, 2011). Other studies have shown that the optimal combination rules for approximating the multi-object pattern can be shifted away from a simple average when objects are correctly positioned for an action between the objects (Baeck et al., 2013) or between an object and a human actor (Baldassano et al., 2017).

Importantly, most objects in natural scenes adhere to meaningful and recurring positional structures: constrained by their function, their relationship with other objects, and physical properties of the world, objects appear in predictable locations, viewpoints, and sizes relative to each other (e.g., sofas facing TV sets, or cars stopping in front of traffic lights). Previous research has demonstrated that the presence of such inter-object regularities facilitates behavioral performance in capacity-limited tasks (Bar, 2004; Chun, 2000; Kaiser et al., 2014a; Oliva and Torralba, 2007; Wolfe et al., 2011), raising the question of how regularly positioned object arrangements are processed in the brain. In the current study, we provide evidence that the visual system integrates representations of regularly positioned objects.

In two fMRI experiments, participants viewed pairs of objects that formed minimalistic versions of scenes (e.g., a "living room", consisting of a sofa and a TV; Fig. 1, **A**). These pairs could be arranged in a regular, typically encountered way (e.g., the sofa facing the TV) or in an irregular way (with the two objects exchanged). Additionally, each single object was presented centrally and in isolation. The multi-voxel response pattern evoked by each pair was then modeled by the average of the patterns evoked by its constituent objects. Replicating previous observations (Kaiser et al., 2014b; MacEvoy and Epstein, 2009, 2011), we find that OSC response patterns evoked by a pair can be accurately predicted by the average response patterns evoked by its constituent objects. Crucially, however, this linear approximation of the pair response depended on the relative position of the objects: Pair approximation in OSC was less accurate for regularly positioned pairs than for irregularly positioned pairs, showing that the independence of multiple object processing breaks down when objects are positioned regularly. More detailed analyses revealed a transformation from independent to integrative representations of regularly positioned object pairs along the posterior-anterior axis of the visual cortex. We interpret this integrative component of multi-object processing as reflecting an emerging tuning to positional structures in real-world scenes.

## 2 Materials and Methods

### 2.1 Participants

Fifteen healthy adults (6 male; mean age 24.6 years, SD=3.3) took part in Experiment 1, and 16 healthy adults took part in Experiment 2 (8 male; mean age 24.7 years, SD=3.0). All participants had normal or corrected-to-normal visual acuity. All procedures were carried out in accordance with the Declaration of Helsinki and were approved by the ethical committee of the University of Trento. Due to excessive head movement, one participant was excluded from Experiment 1. Two participants were excluded from Experiment 2, one due to excessive head movement, and one due to bad functional localizer data, not allowing for a reliable definition of object-selective voxels.

### 2.2 Stimuli

The stimulus set comprised four pairs of objects, which each depicted a minimalistic version of a scene (Fig. 1, **A**). The four scenes were bathroom (toilet and sink), living room (sofa and television), street crossing (car and traffic lights), and playground (seesaw and slide). The stimuli were created by cutting the objects directly from complete scene images, leaving the size and position the same as in the original scene (“regular” condition). For each scene, a second version was created where the object positions were swapped (“irregular” condition). These irregular versions contained the same objects, but the arrangement did not follow the typical arrangement normally experienced in scenes. For each pair, a total of eleven different stimuli were used. In addition to the pair stimuli we also showed the eight single objects centrally, and in isolation. For Experiment 2, additionally a set of 22 scenes (11 indoor rooms, 11 outdoor street scenes) was used; none of these scenes contained any of the objects that constituted the pairs. All stimuli were surrounded by a black frame, which subtended a visual angle of 10° × 7.5°. Stimulus presentation was controlled using MATLAB and Psychtoolbox (Brainard, 1997). The stimulation was back-projected onto a translucent screen placed at the end of the scanner bore. Participants viewed the screen through a tilted mirror mounted to the head coil.

### 2.3 Experiment 1 procedure

The experiment consisted of multiple runs of 5 minutes each. Half of the runs were object pair runs, and half of the runs were single object runs. During object pair runs, the four different pair types (bathroom, living room, street crossing, playground) were shown in each of the two conditions (regular and irregular). During single object runs, all constituent single objects were shown. As a previous study has shown that an approximation of multi-object patterns by single-object patterns is also possible when the positions of the objects do not match (MacEvoy and Epstein, 2011), the single objects were presented centrally. Each run consisted of 18 blocks. Within each block, twelve stimuli were presented, with each stimulus being an exemplar of the same condition (e.g., a bathroom/irregular block would contain only bathroom pairs in irregular configuration). Each block contained all eleven pairs (i.e., exemplars) of a specific condition, and one-back image-level repetition of a specific pair (i.e., exemplar). Each stimulus was shown for 400ms, with an inter-stimulus-interval of 900ms (Fig. 1, **B**); each block thus lasted 15.6s. The first nine blocks of each run consisted of eight stimulation blocks and one block of fixation baseline (in randomized order). The second nine blocks of a run were the same blocks in mirror-reversed order. The experiment comprised ten runs (two participants only completed eight runs), always starting with a pair run and then alternating between single object and pair runs. Participants had to report one-back image repetitions by pressing a button.

**Fig. 1.**
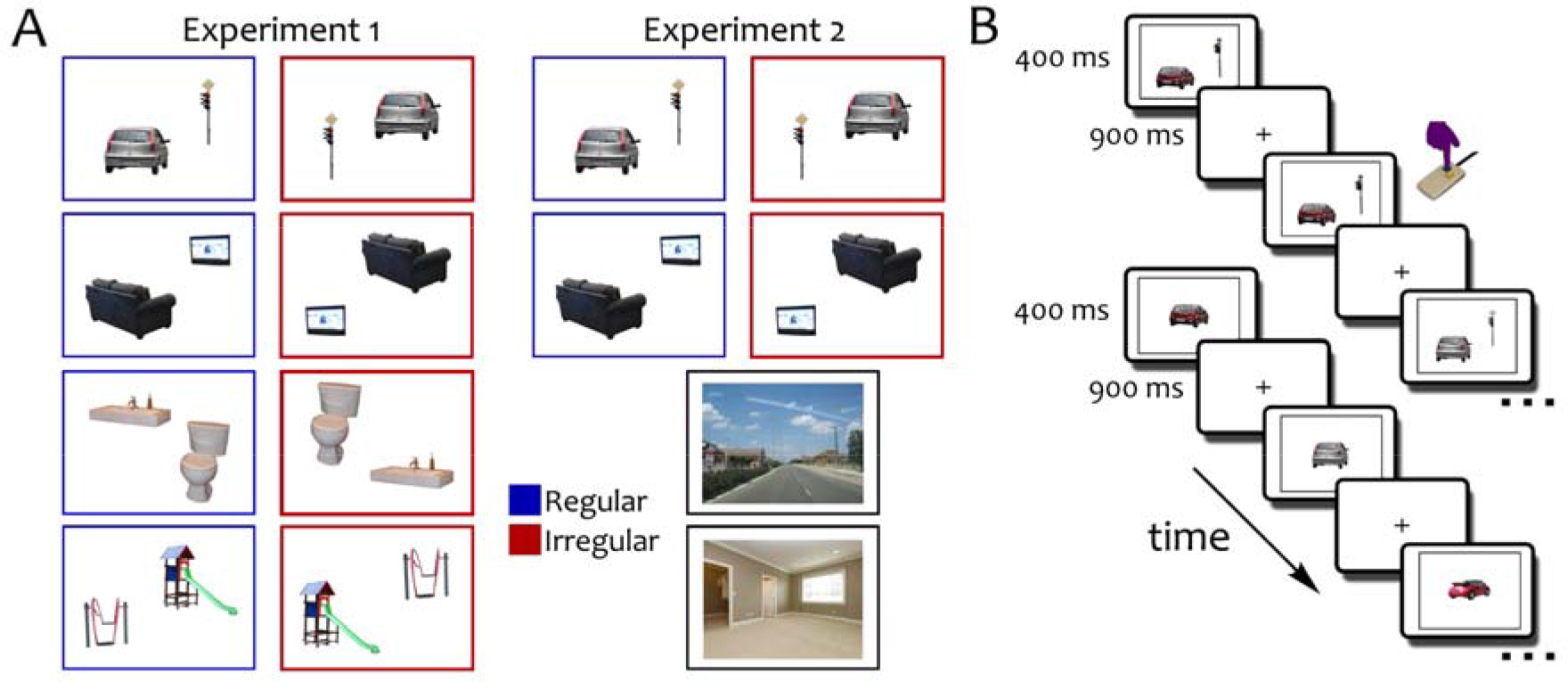
Stimuli and paradigm. **A**, For Experiment 1, four pairs of objects were collected by directly cropping them from scenes of four types (bathroom, living room, street crossing, playground). Object pairs could be positioned in a regular way (as they appeared in the scenes) or in an irregular way (with their positions exchanged). In Experiment 2, only the living room and street crossing pairs were used and, in separate blocks, participants saw images of natural scene images (indoor room versus outdoor street scenes). **B**, Stimuli were presented for 400ms, separated by inter-stimulus intervals of 900ms. In each stimulation block, only pairs from one category and one regularity condition appeared (e.g., regularly positioned cars and traffic lights). Participants responded to one-back image-level repetitions. Similarly, the single objects belonging to the pairs were presented in separate runs, using the same block design and task.

### 2.4 Experiment 2 procedure

The procedure in Experiment 2 was identical to that of Experiment 1 unless otherwise noted. In Experiment 2, only two of the object pairs were used in the pair runs (living room and street crossing), and only their four constituent objects were used in the single object runs. Additionally, in separate runs, the natural scene stimuli were presented (Fig. 1, **A**). These runs had an identical structure to the pair and single object runs. In alternating order, only indoor rooms or outdoor street scenes appeared in a single block. In total, Experiment 2 consisted of 9 runs (3 pair runs, 3 single object runs, 3 scene runs), always starting with a pair run, followed by a single object run and by a scene run (this order was repeated three times).

### 2.5 Functional localizer

In both experiments, participants additionally completed two functional localizer runs of five minutes each. Participants performed a one-back task while viewing images of faces, houses, everyday objects, and phase-scrambled versions of these objects. Each stimulus category included 36 individual exemplars. Within each run, there were four blocks of each stimulus category and four blocks of fixation baseline, with all blocks lasting 16s. Block order was randomized for the first ten blocks and then mirror-reversed for the remaining ten blocks. Each non-fixation block included two one-back stimulus repetitions.

### 2.6 fMRI data acquisition and preprocessing

MR imaging was conducted using a Bruker BioSpin MedSpec 4T scanner (Bruker BioSpin, Rheinstetten, Germany), equipped with an eight-channel head coil. During the experimental runs, T2*-weighted gradient-echo echo-planar images (EPI) were collected (repetition time TR=2.0s, echo time TE=33ms, 73° flip angle, 3 × 3 × 3mm voxel size, 1mm gap, 34 slices, 192mm field of view, 64×64 matrix size). Additionally, a Ti-weighted image (MPRAGE; 1 × 1 × imm voxel size) was obtained as a high-resolution anatomical reference. All resulting data were preprocessed using MATLAB and SPM8. EPI volumes were realigned and coregistered to the structural image. Functional volumes collected during the functional localizer runs were additionally smoothed with a 6mm full-width-half-maximum Gaussian kernel.

### 2.7 Region of interest definition

The BOLD-signal of each voxel in each participant in the localizer runs was modeled using one regressor for each stimulus category, and six regressors for the movement parameters obtained from the realignment procedure. Object-selective cortex (OSC) (Malach et al., 1995) was localized using an object>scrambled contrast, thresholded at p<0.001 (uncorrected). Regions for both hemispheres were concatenated to form a bilateral ROI (Experiment 1: mean ROI size 1000 voxels, SE=35; Experiment 2: mean ROI size 1036 voxels, SE=45). Additionally, an early visual cortex (EVC) ROI was defined by inverse-normalizing an anatomical mask (BA17/18; created from WFUpickatlas for SPM) into individual-subject space (Experiment 1: mean ROI size 996 voxels, SE=8; Experiment 2: mean ROI size 969 voxels, SE=9). For both ROIs, an additional voxel selection criterion was adopted: as we expected that optimally pair-selective voxels allow for better approximation of the pair patterns by single-object patterns (Baeck et al., 2013; MacEvoy and Epstein, 2009), the OSC and EVC ROIs were intersected with voxels that could optimally discriminate between the different pairs (irrespective of regularity condition). These voxels were selected from a whole-brain searchlight classification analysis (see below). A-priori, a voxel count of the 100 most pair-selective voxels was set for all analyses; to rule out that the results were specific to this particular number of selected voxels, the analysis was repeated with different voxel counts ranging from 20 to 500 (see Supplementary Information). For results obtained with a different selection method and for results in scene-selective regions see Supplementary Information.

### 2.8 Univariate Analysis

The BOLD-signal of each voxel in each participant was modeled using one regressor for each stimulus type, and six regressors for the movement parameters obtained from the realignment procedure. Activation differences were computed by averaging beta-weight estimates across voxels for each ROI. Significant differences between the regularly and irregularly positioned pairs were assessed using t-tests.

### 2.9 Multivariate pattern analysis

Multivariate pattern analysis (MVPA) was carried out on a TR-based level using the C0SMoMVPA toolbox (Oosterhof et al., 2016). For each voxel belonging to a specific ROI, TRs corresponding to the conditions of interest were selected by shifting the voxel-wise time-course of activation by three TRs (to account for the hemodynamic delay). Subsequently, for each run separately, activation values were extracted from the EPI-volumes for each TR coinciding with the onset of a specific condition; the activation values were then normalized by z-scoring the values for each voxel. For the decoding analyses, linear discriminant analysis (LDA) classifiers were trained to discriminate the object pairs on all but one runs, and tested on the remaining run; this procedure was repeated so that every run was left out once. Pair classification was also computed in a searchlight decoding analysis (which was used for guiding ROI definition): for this analysis, a spherical neighborhood of 100 voxels was used to identify voxels in visual cortex that could optimally discriminate the different object pairs, irrespective of the regularity condition. Significant decoding was assessed by comparing classifier accuracy to chance performance, using t-tests.

For the pair combination analysis, response patterns were averaged across all TRs belonging to a specific condition, resulting in a single response pattern for every condition. Single-object response patterns for the two objects belonging to a specific pair (e.g., car and traffic light or sofa and TV) were averaged. Thus, for each pair three response patterns were available: (1) the response pattern evoked by the pair presented in the regular arrangement, (2) the response pattern evoked by the pair presented in the irregular arrangement, and (3) the average response pattern evoked by the two constituent objects. To assess pair approximation quality in the regular and irregular conditions separately, the average single object patterns were correlated to their corresponding pair response patterns (within-correlation) and to the pair patterns evoked by the non-corresponding pairs (between-correlation, average of 3 correlations) (Fig. 2).

For the scene approximation analysis in Experiment 2, pair response patterns were correlated with the corresponding scene response patterns (e.g., sofa/TV and indoor room scene; within-correlation) and the non-corresponding scene patterns (e.g., sofa/TV and outdoor street scene; between-correlation), separately for regular and irregular pairs. The difference between these within- and between-correlations was used as a measure of pair approximation quality. Approximation quality was then compared for the regular and irregular pairs by using paired t-tests. All raw correlations were Fisher-transformed prior to statistical analysis.

**Fig. 2.**
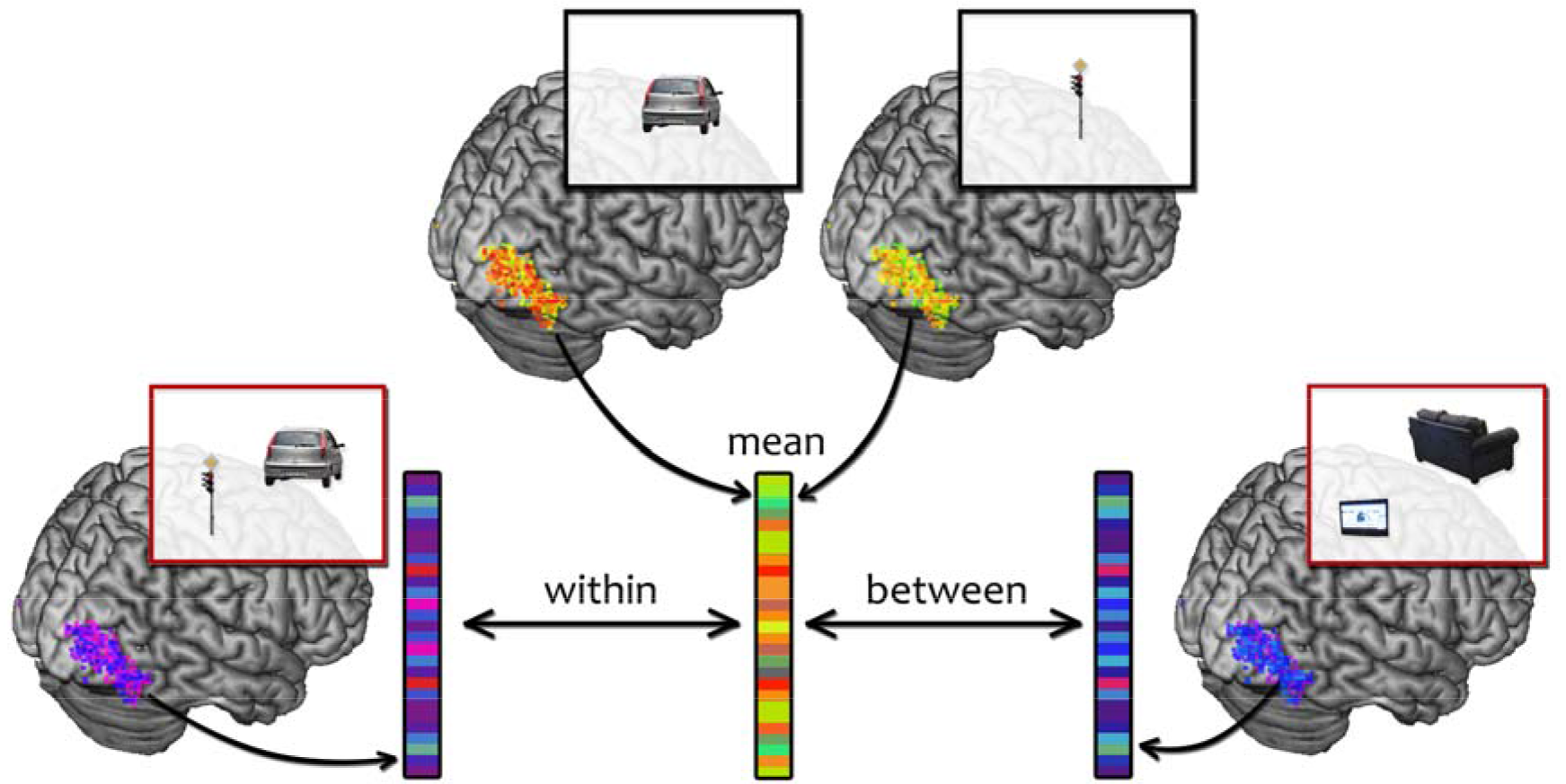
Schematic illustration of the pair approximation analysis. For each ROI, the response pattern across hemispheres was extracted both for the single object and pair conditions. The single object patterns corresponding to each pair were then averaged and correlated with the pair-evoked pattern. The difference between the correlations with corresponding pairs (within-correlation) and non-corresponding pairs (between-correlation) was taken as a measure of approximation quality. This analysis was done separately for the regular and irregular pairs.

### 2.10 Combined searchlight analysis

To characterize the emergence of the regularity effect along the visual processing hierarchy, we performed a searchlight analysis, where we systematically moved a spatial voxel neighborhood along the anterior-posterior axis. For this analysis, the data from both experiments was combined. First, the occipitotemporal cortex (defined based on an anatomical mask) was divided into 40 slices. Each slice spanned 10mm in the anterior-posterior (Y) direction, with an inter-slice distance of 2mm (thus, individual slices were overlapping). X and Z coordinates of the occipitotemporal region were fully covered, with the exception that the medial parts of occipital cortex were masked out to avoid sampling of EVC voxels. The 40 slices were then inverse-normalized to individual-participant space. Separately for each participant, within each slice the 100 voxels that allowed the most accurate pair decoding in a searchlight analysis were selected (similarly to the OSC analyses, see above). Subsequently, the pair approximation analysis (see above) was performed for these 100 voxels within each slice. To assess the general object-selectivity of the voxels in each slice, we computed the difference between the GLM beta weights for intact and scrambled object in the functional localizer runs (see above). To identify slices yielding significant effects, we used a threshold-free cluster-enhancement procedure (Smith and Nichols, 2009) with default parameters, using multiple-comparison correction based on a sign-permutation test (with null distributions created from 10,000 bootstrapping iterations) as implemented in CoSMoMVPA (Oosterhof et al., 2016). The resulting Z-values were thresholded at Z>1.96 (i.e., p<0.05).

## 3 Results

### 3.1 Experiment 1

To investigate whether positional regularities impact neural responses in object-selective visual cortex (OSC), we presented participants with pairs of objects that commonly appear together in real world scenes, such as in a bathroom (toilet and sink), in a living room (sofa and TV), on a street crossing (car and traffic lights), or at a playground (seesaw and slide). These objects were directly cut from natural scene images and presented on a blank background (“regular” condition; Fig. 1, **A**). To investigate whether the way in which these objects typically co-occur in the real world influences neural responses, we added a condition where we reversed the object locations (“irregular” condition, Fig. 1, **A**), keeping everything else identical. Additionally, the single objects belonging to the pairs were presented centrally (Fig. 1, **B**).

Functional MRI analyses primarily focused on two regions of interest: functionally defined object-selective cortex (OSC) and anatomically defined early-visual cortex (EVC) (see Supplementary Information for results in scene-selective ROIs). Based on previous work (Baeck et al., 2013; MacEvoy and Epstein, 2009), we expected stronger effects in areas of OSC that preferentially represent information about the object pairs. To define voxels that contribute most to the coding of the object pairs, a whole-brain searchlight analysis was performed (see Materials and Methods), where linear classifiers were used to discriminate the four object pairs, regardless of their positioning (i.e., collapsed across the regular and irregular pairs). Based on this searchlight analysis, within OSC and EVC the 100 voxels that contributed most to the classification of the four pairs were selected (Fig. 3, **A**). Other voxel counts and voxel selection methods gave similar results (see Supplementary Information).

**Fig. 3.**
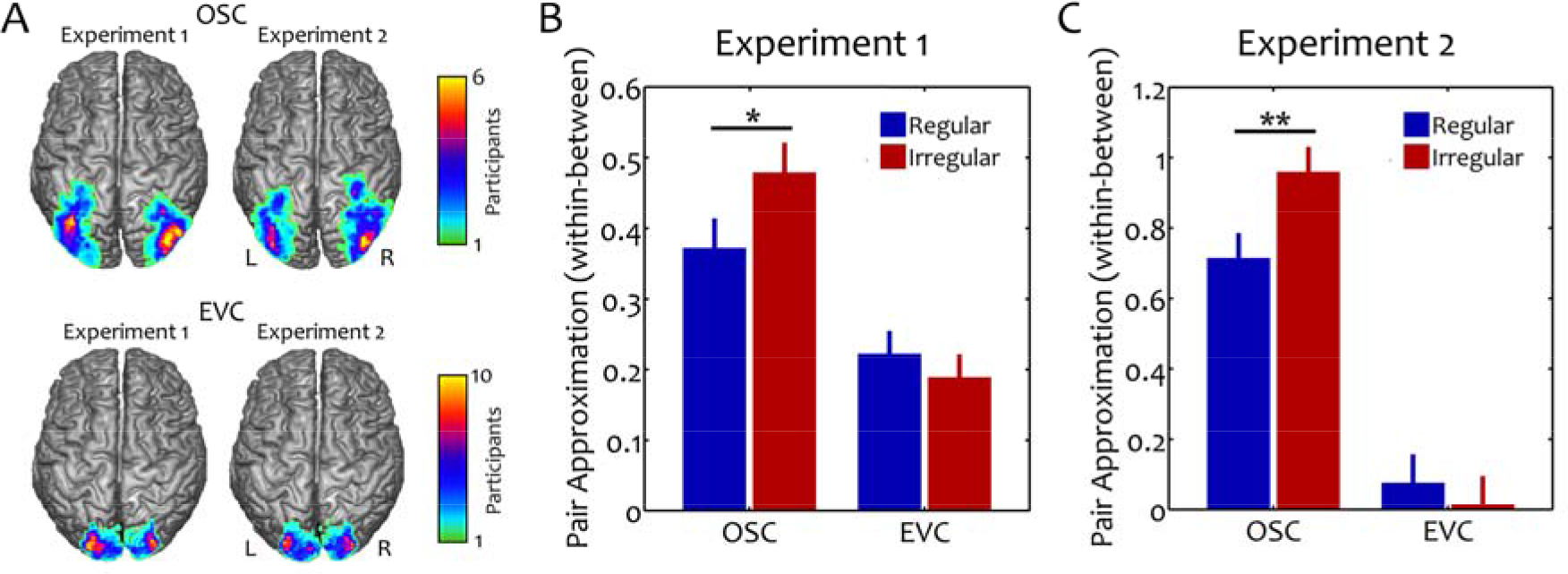
Pair approximation results. **A**, Object selective cortex (OSC) and early visual cortex (EVC) maps depicting voxel-wise overlap between participants: Regions were defined on a single-subject level by selecting the 100 voxels that best discriminated the four pairs across the regularity conditions (see Materials and Methods) within either a functional OSC mask or an anatomical EVC mask. For illustration purposes, the regions were normalized into standard space and overlaid on a brain template using MRIcron (Rorden and Brett, 2000). **B**, The pair approximation analysis for Experiment 1 revealed a more accurate approximation of the irregularly positioned pairs than of the regularly positioned pairs in OSC, while no effect was found in EVC. **C**, In Experiment 2, this pattern of results was replicated, demonstrating sensitivity for object regularity structure in OSC, but not EVC. All error bars reflect standard errors of pairwise differences. (*p<0.05, **p<0.01)

First, we wanted to confirm that the four object pairs were reliably discriminable in both OSC and EVC. Separately for regular and irregular pairs, a multivariate classification analysis of response patterns across voxels was performed. In a leave-one-run-out fashion, linear discriminant analysis (LDA) classifiers were used to classify response patterns to the four different pairs (see Materials and Methods). In both OSC and EVC, these classifiers performed highly above chance level (OSC: 42%; EVC: 40%), as expected based on the voxel selection method. For both regions, no reliable difference in decoding accuracy was found when comparing the regular and irregular conditions directly (OSC: t[13]=0.8, p=0.41; EVC: t[13]=1.2, p=0.25). Similarly, no difference in univariate activation values between the two conditions was observed (OSC: t[13]=0.7, p=0.48; EVC: t[13]=1.1, p=0.29). This pattern of results argues against overall differences in signal quality, for example due to differences in attentional engagement for the two conditions: such an effect would be expected to result in a univariate and/or classification difference between the regular and irregular conditions.

Next, we tested whether the average of the response patterns evoked by the single objects could more accurately approximate the pattern evoked by the corresponding pairs than the pattern evoked by the non-corresponding pairs: for example, is the average of the response patterns for cars and traffic lights more similar to the response pattern for the car and traffic light pair than to the other pairs (e.g., to a TV and a sofa)? Two types of correlations were computed: within-correlations, of the average single object response patterns and the corresponding pair response patterns; and between-correlations, of the average single object response patterns and the other, non-corresponding pair response patterns (see Materials and Methods; Fig. 2). For both regular and irregular pairs separately, the difference between these within- and between-correlations was taken as a measure of pair approximation quality.

In OSC, within-correlations were significantly higher than between-correlations, both for regular and irregular pairs, indicating that the mean of the constituent object responses offered a reliable approximation of the pair responses (regular: t[13]=6.33, p<0.001; irregular: t[13]=11.2, p<0.001; Fig. 3, **B**). Similar effects were observed in EVC (regular: t[13]=6.1, p<0.001; irregular: t[13]=4.7, p<0.001). Next, to investigate whether positional regularity affected this approximation, we directly compared regular and irregular pair approximation. Crucially, pair approximation was better for irregular pairs than for regular pairs (t[13]=2.5, p=0.02). By contrast, no such difference was found in EVC (t[13]=1.0, p=0.32; interaction with region: t[13]=2.9, p=0.01). These findings indicate that multiple object processing in OSC is sensitive to typical real-world positioning.

### 3.2 Experiment 2

In a second fMRI experiment we aimed to replicate the pattern of results. The structure of the experiment was largely identical to Experiment 1 (see Materials and Methods) and participants viewed images from two of the pairs also used in Experiment 1 (sofa and TV; car and traffic lights).

First, as in Experiment 1, discriminability of the response patterns for the regular and irregular pairs was assessed using a two-way linear classification analysis in OSC and EVC. In both regions, classifiers performed highly above chance level (OSC: 69%; EVC: 64%) and no difference in decoding accuracy was found when comparing the regular and irregular conditions directly (OSC: t[13]=0.02, p=0.99; EVC: t[13]=0.7, p=0.51). Additionally, no difference was observed in univariate activation values for the two conditions (OSC: t[13]=0.01, p=0.99; EVC: t[13]=1.0, p=0.32).

Second, separately for regular and irregular pairs, constituent single-object response patterns were averaged for each pair to test how well these average response patterns could approximate the two pairs. As in Experiment 1, in OSC the average of the response patterns evoked by the single objects was more similar to the pattern evoked by the corresponding than by the non-corresponding pairs, both for the regular pairs (t[13]=11.1, p<0.001) and the irregular pairs (t[13]=11.7, p<0.001; Fig. 3, **C**). By contrast (and unlike Experiment 1), the average single object patterns did not accurately approximate the pair patterns in EVC (regular pairs: t[13]=0.8, p=0.44; irregular pairs: t[13]=0.2, p=0.87). Most importantly, Experiment 2 replicated the key finding of Experiment 1: a difference between the approximation of regular and irregular pairs in OSC (t[13]=3.4, p=0.004), with no such difference in EVC (t[13]=0.8, p=0.46; interaction across regions: t[13]=3.3, p=0.006). These results closely resemble the pattern of results obtained in Experiment 1, providing further evidence for an influence of positional regularities on multi-object coding in OSC.

Experiment 2 was additionally designed to test a possible explanation for the regularity effect: perhaps regularly positioned object pairs more strongly induced a mental representation of the corresponding scene (e.g., the spatial layout of the scene), thereby distorting the representation of the individual objects. To test this account, participants viewed images of room scenes and street scenes during separate blocks of Experiment 2 (Fig. 1, **A**). None of these scenes contained any of the objects belonging to the pairs. OSC response patterns to these scenes were reliably discriminable in a two-way classification analysis (mean accuracy: 64%; t[13]=9.1, p<0.001). To determine whether the relatively inaccurate pair approximation for regularly positioned pairs in OSC reflected the activation of the corresponding scene representations (e.g., living room for sofa-TV pairs), we correlated the activation patterns evoked by the pair stimuli to the activation patterns of the scene stimuli. Separately for the regular and irregular conditions, this correlation was computed within scene type (i.e., sofa/TV and indoor room scene, and car/lights and outdoor street scene) and between scene type (i.e., sofa/TV and outdoor street scene, and car/lights and indoor room scene). The difference of these within- and between-correlations was taken as a measure of scene approximation quality. In OSC, this scene approximation was not significantly different from zero, both for regularly (t[13]=0.4, p=0.69) and irregularly (t[13]=0.2, p=0.87) positioned pairs and no difference was found when directly comparing discriminability for regular and irregular pairs (t[13]=0.7, p=0.50). Similar results were obtained in scene-selective regions of visual cortex (see Supplementary Information). These results suggest that the differential coding of regular and irregular pairs in OSC does not reflect a difference in the degree to which they activate scene category response patterns.

### 3.3 Transformation from independent to integrative coding

The results from Experiments 1 and 2 demonstrate that a linear combination of single object response patterns accurately predicts the response patterns evoked by both regularly and irregularly positioned pairs. However, a relative disruption of this approximation was observed in the regular condition. This pattern of results suggests that responses to regularly positioned pairs reflect both an independent processing component and an integrative component that disrupts this independent processing to some degree. As our region of interest analysis only focused on two larger portions of the visual cortex (EVC versus OSC), it could not reveal which parts of OSC are housing the independent and integrative components of multi-object processing. To reveal the transformation from independent to integrative processing at finer anatomical resolution we tracked the overall approximation quality (reflecting the independent component of multi-object representations) and the regularity effect (reflecting the relative disruption of independent coding in the regular condition) along the visual hierarchy. Combining the data from both experiments, we performed a searchlight analysis along the posterior-anterior axis of the occipitotemporal cortex (see Materials and Methods). In brief, anatomically defined occipitotemporal cortex was divided into slices (width: 10mm) varying along the y-coordinate (in MNI space) in steps of 2mm (see Fig. 4, **A**). For each of these slices, comparably to the OSC analyses for Experiments 1 and 2, the 100 best pair-representing voxels were selected based on individual-participant searchlight maps. Then, for each slice separately, the pair approximation analysis was performed (as illustrated in Fig 2), leading to a pair approximation value (within-correlation vs. between-correlation) and a regularity effect for each slice (Fig. 4, **B**). Generally, approximation quality across the two regularity conditions (i.e., the mean of the two conditions) was very robust and significantly above chance for all slices centered between Y=−100 and Y=−34 (Fig 4, **C**). To detect regularity effects, the difference of the linear approximation quality in the regular and irregular conditions was compared to zero. This analysis revealed an effect that emerged between Y=−80 and Y=−84, and later between Y=−64 and Y=−38 (Fig 4, **D**). This pattern of results shows that highly accurate linear approximation can be observed at earlier processing stages than the relative breakdown of this approximation in the regular condition, suggesting that the integrative multi-object coding for regularly positioned objects is preceded by an independent representational stage.

**Fig. 4.**
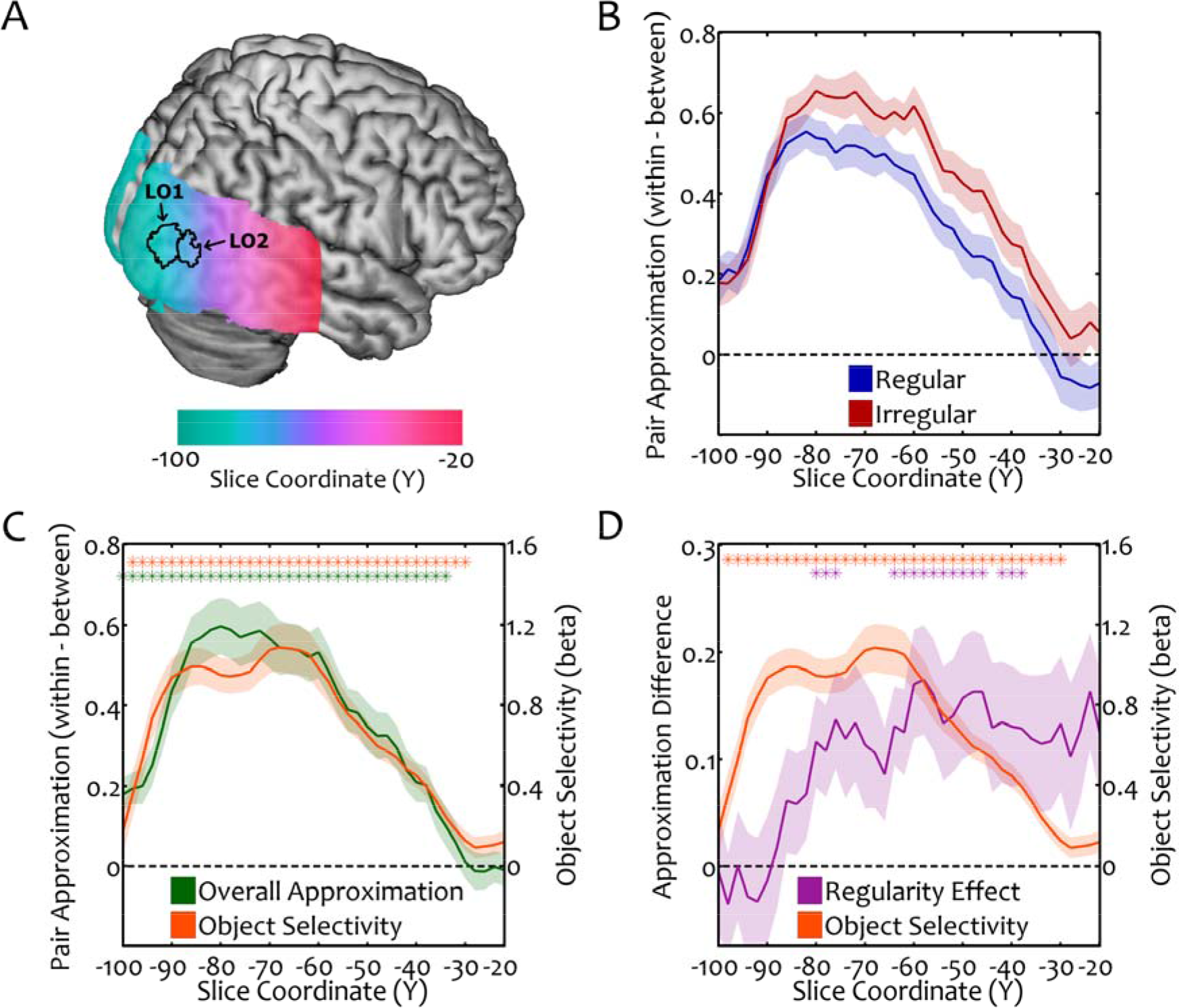
Combined searchlight analysis. **A**, Searchlight analysis was performed for slices of occipitotemporal cortex (iomm width) with varying y-coordinate, in steps of 2mm. Slices were defined in standard space (based on IVIN I coordinates) and back-projected into individual subject space. Subsequently, within each slice, the 100 best pair-representing voxels were selected on an individual-participant level. **B**, Pair approximation (expressed as the difference between within- and between correlations) for regular and irregular pairs in slices along the y-coordinate. **C**, Overall approximation (i.e., the mean of the regular and irregular conditions) was generally very accurate throughout the visual hierarchy, and showed a striking correspondence with the degree of object-selectivity. **D**, Differences between the regular and irregular conditions emerged starting from Y=−80. In contrast to the overall approximation, the regularity effects along the hierarchy did not correspond well with the degree of object-selectivity. (*p<0.05, corrected for multiple comparisons). The transition from independent to integrative coding might occur between object-selective regions LO1 and LO2 (**A**, region maps taken from Wang et al., 2015), with independent coding in LO1 and additional integrative coding in LO2 (see Discussion).

To assess how the overall approximation quality and the regularity effect relate to object selectivity, the difference of the intact and scrambled object conditions in the functional localizer was computed for each slice (see Materials and Methods). All slices between Y=−98 and Y=−30 exhibited significant object-selectivity (Fig. 4, **C/D**). Interestingly, across slices object-selectivity almost perfectly predicted overall approximation quality (r=0.97, p<0.001). By contrast, object-selectivity did not significantly predict the regularity effect across slices (r=−0.05, p=0.75). These results suggest that the independent component of multi-object processing is tightly linked to the degree of object-selectivity, while the influence of object regularity emerges later in the visual hierarchy. In sum, this analysis shows that overall linear approximation (reflecting the independent component of multi-object processing) and its relative breakdown (reflecting an effect of positional structure) can be anatomically dissociated, revealing a transformation from independent to integrative coding of multi-object displays along the visual processing hierarchy.

## 4 Discussion

In two fMRI experiments, we demonstrate non-independence in the visual cortex representation of multi-object displays, revealing a transformation from independent to integrative representations along the visual processing hierarchy. Response patterns in OSC evoked by regularly positioned object pairs were less accurately modeled by the response patterns evoked by their constituent objects. The relative breakdown of this linear response approximation reflects a disruption of independent multi-object coding when objects adhere to their typical real-world positions. The regularity effect was significant in functionally localized OSC, but not in EVC. (Mixed results were obtained in scene-selective regions; see Supplementary Information.) However, further analyses revealed that object selectivity was closely related, along the posterior-anterior axis, to independent object coding rather than integrative object coding, which emerged later in the visual hierarchy.

The searchlight analysis along the posterior-anterior axis of the occipitotemporal cortex suggests that the coding of multiple, regularly positioned objects reflects two dissociable components: an independent processing component and an integrative component. Independent processing can be observed starting at early stages of the visual object processing hierarchy, and is tightly linked to object selectivity. The striking correspondence between object-selectivity and pair approximation suggests that the same neural mechanisms support single-object coding and (independent) multi-object coding. By contrast, at later stages of the processing hierarchy, pair approximation quality diverges for regular and irregular pairs, reflecting additional, integrative processing for regularly positioned pairs that (partly) disrupts independent coding. Interestingly, this relative disruption cannot be explained by differences in object-selectivity along the visual hierarchy.

We speculate that the independent and integrative components of multi-object coding reflect activity in different sub-regions of object-selective cortex, with a more posterior sub-region – possibly LO1 (Larsson and Heeger, 2006) (Y=−90, SE=5) − primarily reflecting independent coding and a more anterior sub-region – possibly LO2 (Larsson and Heeger, 2006) (Y=−83, SE=6) – also containing an integrative component. This proposal is supported in our current data: Repeating the main pair approximation analysis using probabilistic LO1/LO2 group masks [^1^ ROIs were generated by inverse-normalizing the group masks into individual-subject space. To ensure sufficient voxel counts for the analysis, all voxels exceeding a probability of 25% of belonging to either region were selected. Overlapping voxels were grouped into the higher-probability region.] (Wang et al., 2015; Fig. 4, **A**), we found that pair approximation was disrupted in LO2 (t[27]=2.0, p=0.05), but not in LO1 (t[27]=0.7, p=0.47; interaction across regions: t[27]=2.3, p=0.03). This difference in integrative object processing is consistent with a previously reported difference in spatial summation along the posterior-anterior axis, with a higher rate of information compression in the more anterior LO2 (Kay et al., 2013). However, further studies employing retinotopic mapping techniques are needed to more precisely pinpoint the effects to specific sub-regions of object-selective occipitotemporal cortex.

The disruption of independent coding observed here is unlikely to reflect differential semantic associations between objects (e.g., that car and traffic light are semantically related). Previous electrophysiological studies have demonstrated that such associations between objects can alter neural tuning properties in visual cortex: The associative pairing of two objects can shape neural responses in a way that neurons become selective to both of the individual objects (Messinger et al., 2001; Sakai et al., 1991). However, in our study the objects were equally related in their semantic content in the regular and irregular conditions – the crucial difference was the positional structure. An alternative explanation is that our results reflect the activation of object group representations that are shaped by the co-occurrence structure of objects in real-world environments. Such group representations have been proposed for the cortical representation of people, where information from the face and body is integrated into a person representation (Bernstein et al., 2014; Harry et al., 2016; Schmalzl et al., 2012; but see Kaiser et al., 2014b). The additional recruitment of group representations would disrupt the linear approximation of multi-object patterns: if neurons mediating the group representation are distributed differently across voxels from neurons mediating the individual object representations, the pair patterns suffers a distortion (relative to the average of the individual object patterns). Further studies are required to probe the nature of these group representations at a finer grained level.

A number of previous studies have explored relational coding of objects in the context of action relationships. Some of these studies have reported univariate activation differences between objects positioned correctly or incorrectly for performing a specific action (Kim and Biederman, 2011; Kim et al., 2011; Roberts and Humphreys, 2010). Others have used similar methods as the current study to demonstrate that action relationships alter combination rules in the processing of multiple objects (Baeck et al., 2013; Baldassano et al., 2017). Notably, the regularity effects observed in these studies have been attributed to processing asymmetries favoring the “active” object (e.g., a hammer) over the “passive” object (e.g., a nail) in a pair (Baeck et al., 2013; Riddoch et al., 2003). By contrast, our approach focused on real-world object structure more generally, avoiding such direct functional interactions of the two objects. Furthermore, in our study the positional structure of the object pairs was solely manipulated by exchanging the position of the single objects, without introducing changes of low-level visual characteristics (like physically connecting the objects differently; see Kim and Biederman, 2011; Baldassano et al., 2017). Therefore, the relative disruption of independent coding in the regular condition observed in our data cannot be explained by low-level visual interactions between the two stimuli. Our results thus provide strong evidence for sensitivity for positional structure in visual cortex, which is not attributable to a shift of weights in favor of one of the objects and is not due to low-level grouping.

Previous studies using similar paradigms have often shown single objects and multiple objects in identical, peripheral locations (Baeck et al., 2013; Kaiser et al., 2014b; MacEvoy and Epstein, 2009). By contrast, our study employed a design where the single objects were presented centrally and thus in different retinotopic locations as within the pairs (MacEvoy and Epstein, 2011). In this case, the successful approximation of the pair responses – and thus the breaking down of this approximation – must emerge from neural populations with sufficiently large receptive fields and sufficient location tolerance (this also becomes apparent in the higher pair approximation quality in OSC, as compared to EVC). In natural environments, this location tolerance might be a crucial property of multiple object representations: In the real world, objects frequently change retinotopic position but their relative position is more stable. Object group representations may thus help to maintain a coherent and stable representation of a scene.

How does the neural sensitivity for positional regularities arise from the typical relative positioning of objects? In the present study, we interchanged the position of the object pairs to disrupt positional structure while keeping the individual objects identical. In the irregular displays, positional structure was disrupted in multiple ways: The objects did not adhere to their typical relative viewpoints and sizes, the laws of physics were partly violated, and the typical functional relationships between the objects were disrupted. All these factors constitute properties of positional object structures in real-world scenes, so that all of them potentially contribute to the effects observed here. Thus, while our study provides clear evidence for a role of positional structure in multi-object representations, further studies are needed to disentangle the contributions of the different aspects our manipulation entailed.

The sensitivity for positional structure observed here may reflect the grouping (or integration) of multiple, regularly positioned objects. Such grouping may provide an efficient way to deal with the large number of objects contained in natural scenes (Kaiser et al., 2014a). As a result of limitations in visual capacity, parallel processing of objects is associated with competition for representation between individual stimuli: When multiple objects need to be processed at the same time, neural processing becomes less efficient and behavioral performance decreases (Desimone and Duncan, 1995; Franconeri et al., 2013; Kastner and Ungerleider, 2001). Processing objects in meaningful chunks might help to reduce the complexity of a scene by reducing the number of competing objects. Behavioral evidence suggests that regularly positioned objects can be more efficiently processed, as measured in various visual tasks (Gronau and Shachar, 2014, 2015; Kaiser et al., 2014a, 2015; Stein et al., 2015). These behavioral benefits have been linked to a reduction in competitive interactions in visual cortex when objects follow positional regularities (Kaiser et al., 2014a). Our current results provide an explanation for this reduction of neural competition: Regularly positioned arrangements of objects are not recruiting independent and competing object representations, but are to some degree integrated into group representations at higher stages of visual cortex. Such group representations may constitute an integral step in the visual processing hierarchy, bridging the gap between individual object coding and the representation of whole scenes.

## Acknowledgements

The research was supported by the Autonomous Province of Trento, Call "Grandi Progetti 2012", project "Characterizing and improving brain mechanisms of attention – ATTEND". This project has received funding from the European Research Council (ERC) under the European Union’s Horizon 2020 research and innovation programme (grant agreement No 725970).

